# Potential Role of Red Palm Oil Supplemented Diet on Oxidative Stress Enzymes in *Plasmodium Berghei* Induced Malaria

**DOI:** 10.1101/2021.03.18.435769

**Authors:** Temitope Daniel Adeleke, Olawale Abiodun Adejumobi, Franklin Folasele Akinola, Oluwatosin Abidemi Salau, Oyeronke Suebat Uthman-Izobo

## Abstract

**Background:** Malaria parasites are very vulnerable to oxidative stress during the part of their life cycle when they inhabit the erythrocytes. Studies have shown that dietary intake of antioxidant plays a role in stabilizing oxidative stress.

**Methods:** The objective of this research work was to examine the antioxidative effect of red palm oil on *Plasmodium berghei* malaria induced oxidative stress. Sixty (60) mice were distributed into five groups. Group A served as the negative control (healthy mice with normal feed); group B as positive control (healthy mice fed with red palm oil without malaria parasite.while the other groups (C to E) served as the test groups. Group C served as group of healthy mice fed with red palm oil (pelletized), infected with malaria parasite without antimalaria drug. Group D served as group of healthy mice fed with red palm oil (pelletized), infected with malaria parasite and treated with amodiaquine. Group E served as group of healthy mice fed with normal feed, infected with malaria parasite and treated with amodiaquine. The parasitemia levels were estimated on days 1,4 and 5. The activity of oxidative stress enzymes biomarkers were determined spectrophotometrically.

**Result:** Group A showed a statistically significant increase in the activity of SOD (1.90 ± 0.16 units/mg protein), GST (1.68 ± 0.086 units/L) compared to group C, SOD (3.54 ± 0.83 units/mg protein), GST (2.12 ± 0.20 units/L). Group B showed a statistical significant decrease in the activities of SOD (3.22 ± 0.33 units/mg protein), Catalase (49.11 ± 2.35 µmol/min), GSH-R (31.50 ± 2.48 units/L) compared to group E, SOD (2.18 ± 0.39 units/mg protein), Catalase (44.07 ± 3.88 µmol/min), GSH-R (27.75 ± 1.64 units/L).

**Conclusion:** The dietary intake of red palm oil helps to reduce free radical mediated injury to the tissue thus preventing oxidative stress induced by malaria or any other factors.

## Introduction

Malaria is an infectious disease caused by protozoan parasites belonging to the genus *Plasmodium* and phylum *Apicomplexa*[1]. Four Plasmodium species have been well known to cause human malaria, namely, *P. falciparum, P. vivax, P. ovale, and P. malariae*[2]. Malaria currently accounts for about 200 million infections each year, with more than 500,000 deaths annually and is estimated to kill one child every 30 seconds [3]. The most common symptoms of malaria include; headaches,body aches, fever, convulsions, vomiting, dehydration, loss of appetite, weight loss, lethargy, low blood levels, coughing, sneezing, fast breathing, restlessness, and depression [4]. Malaria is associated with increased production of free radicals whose activities can be reduced by antioxidants[5]. It has also been documented that malaria parasites exert oxidative stress within parasitized red blood cells[6]. Biochemically, some of the symptoms are believed to have been caused by the induction of oxidative stress. Meanwhile, oxidative stress is defined as an imbalance between oxidants and antioxidants in favour of the oxidants, potentially leading to damage in the body. Oxidants are formed as a normal product of aerobic metabolism but can be produced at elevated rates under pathophysiological conditions [7]. However, oxidative stress will increase continuously when the antioxidants in circulation are unable to reduce the oxidants present in the body. Furthermore, oxidative stress markers in infected humans and rats are found in high levels compared to uninfected controls. In such cases, oxidative stress seems to result from increased production of free radicals, a fact suggested by increased malondialdehyde (MDA), an important lipid peroxidation marker, and not from a decrease in levels of antioxidants, reinforcing the suggestion that oxidative stress is an important mechanism in parasite infection [8]. Antioxidants are readily available in diet and form part of food condiments. Daily diet rich in antioxidant foods include; red palm oil [9]. Red palm oil is loaded with so many nutrients and antioxidants. Itis a natural food/soup condiment in the West African countries. The high antioxidant content of the oil quenches free radicals and keeps inflammation under control[10]. Red palm oil, as the second largest consumed vegetable oil in the world is obtained from a tropical plant*, Elaesisguineensis* contains 50 percent saturated fatty acids, and it contains a high amount of the antioxidants, beta-carotene, and vitamin E [11]. A study had suggested that red palm oil could minimize oxidative damage through its potential ability to increase antioxidant enzymes and it may therefore play a role in the prevention and treatment of oxidative injuries to cells [12]. In this experimental research study, *Plasmodium berghei,* a species of *plasmodium* known to be infectious to rodents (mice) has been used as a model. This is because it is a valuable model organism for the investigation of human malaria as the organisms are similar in most essential aspects of morphology, physiology and life cycle of this parasite is simple and safe [13].

## Materials and Methods

### Study site

Thisstudy was conducted at the Institute for Advance Medical Research and Training (IMRAT), College of Medicine, UCH, Ibadan, Oyo State, Nigeria

### Study design

The study is an experimental study.

### Animal Care

Forty mice of mixed sex with average weight of 19g obtained from an animal breeder were used for this study. All experimental protocols were duly observed according to the guidelines governing the care and use of experimental animals. All animals were made to receive humane care in accordance with the principle of laboratory animal care of the National Society of Medical Research. Animals were housed and acclimatized in the departmental animal room for a week, assigned into five groups [Table 1]. Each group had twelve (12) animals, labelled with their respective group letters with picric acid and, the experimental and the positive control groups were inoculated with *Plasmodium berghei* using the intravenous route of administration. Amodiaquine was administered as the antimalarial drug. All the animals were given access to water and they were fed for seven days.

**Table 1:**
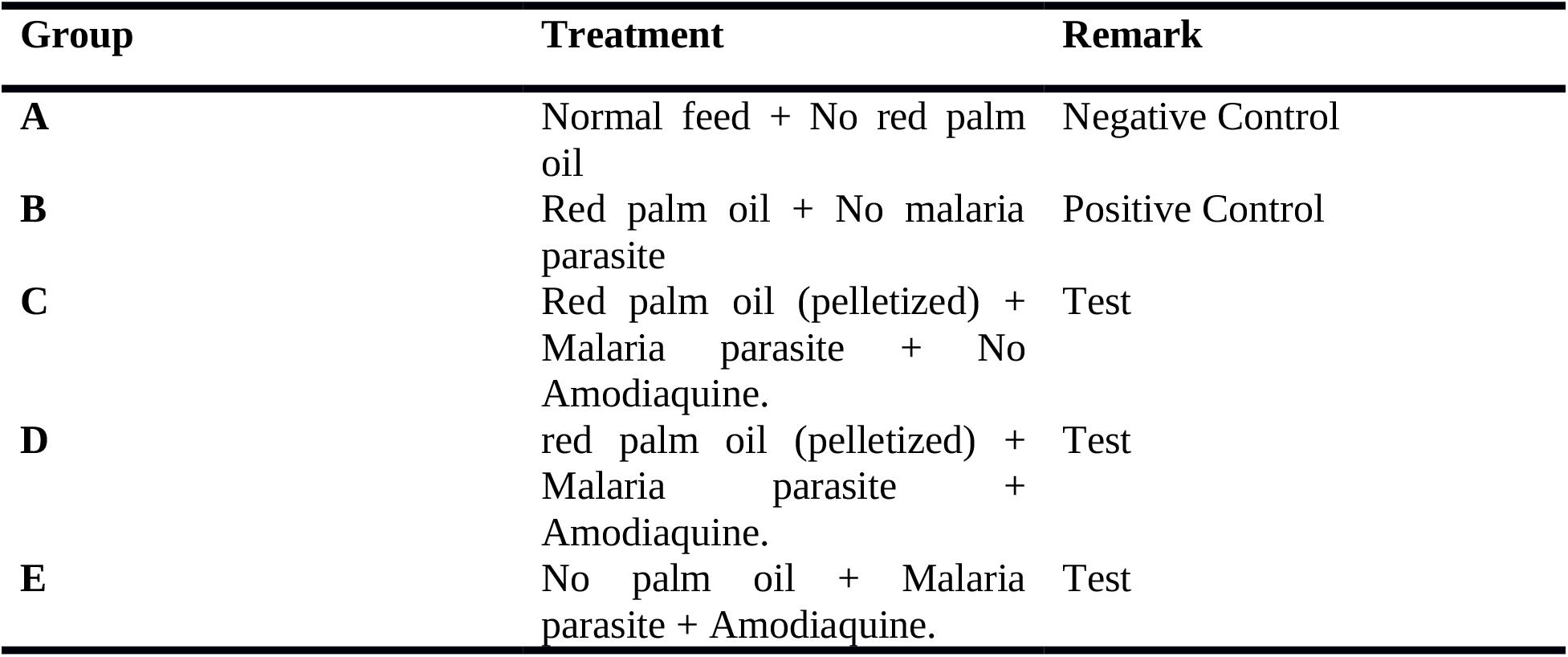
Experimental protocol table.

The study received the approval of the Ethical Committee of Institute for Advance Medical Research and Training (IMRAT), College of Medicine, UCH, Ibadan, Oyo State, Nigeria where the research work was also carried out.

### Blood Collection

Blood was collected from the heart of the mice using 1ml needle and syringe. Immediately after sacrifice, blood was transferred to an eppendorf sample bottle, centrifuged to separate serum and the samples were transported to the laboratory for biochemical analysis. Also, the liver of all the surviving animals were collected into a small capped bottle. The samples were protected from sunlight with aluminum foil during transportation from the animal house to the laboratory for biochemical analysis.

### Serum preparation

The blood collected into the plain bottles was allowed to clot, then centrifuged in 4000 rpm and the supernatant was collected into another plain bottle for analysis

### Determination of catalase (CAT) activity

Catalase activity was determined using direct colorimetric method[14].

### Determination of glutathione reductase (GSH) activity

The method of estimation of the level of reduced glutathione (GSH) determined by [15].

### Determination of superoxide dismutase (SOD) activity

SOD activity was determined by the method [16].

### Assay of glutathione peroxidase (GPx) activity

A proposed method was adopted for assaying the activity of peroxidase [17].

### Assay of glutathione S-Transferase (GST) activity

Glutathione S-transferase was assessed by the method [18].

### Statistical Analysis

Values obtained from the study expressed as mean ± standard deviation were compared using the independent student t-test and significance was measured at *P*<0.05.

## Results

The results of the study are presented in Tables 2 and 3 and Fig. 1.

**Table 2:**
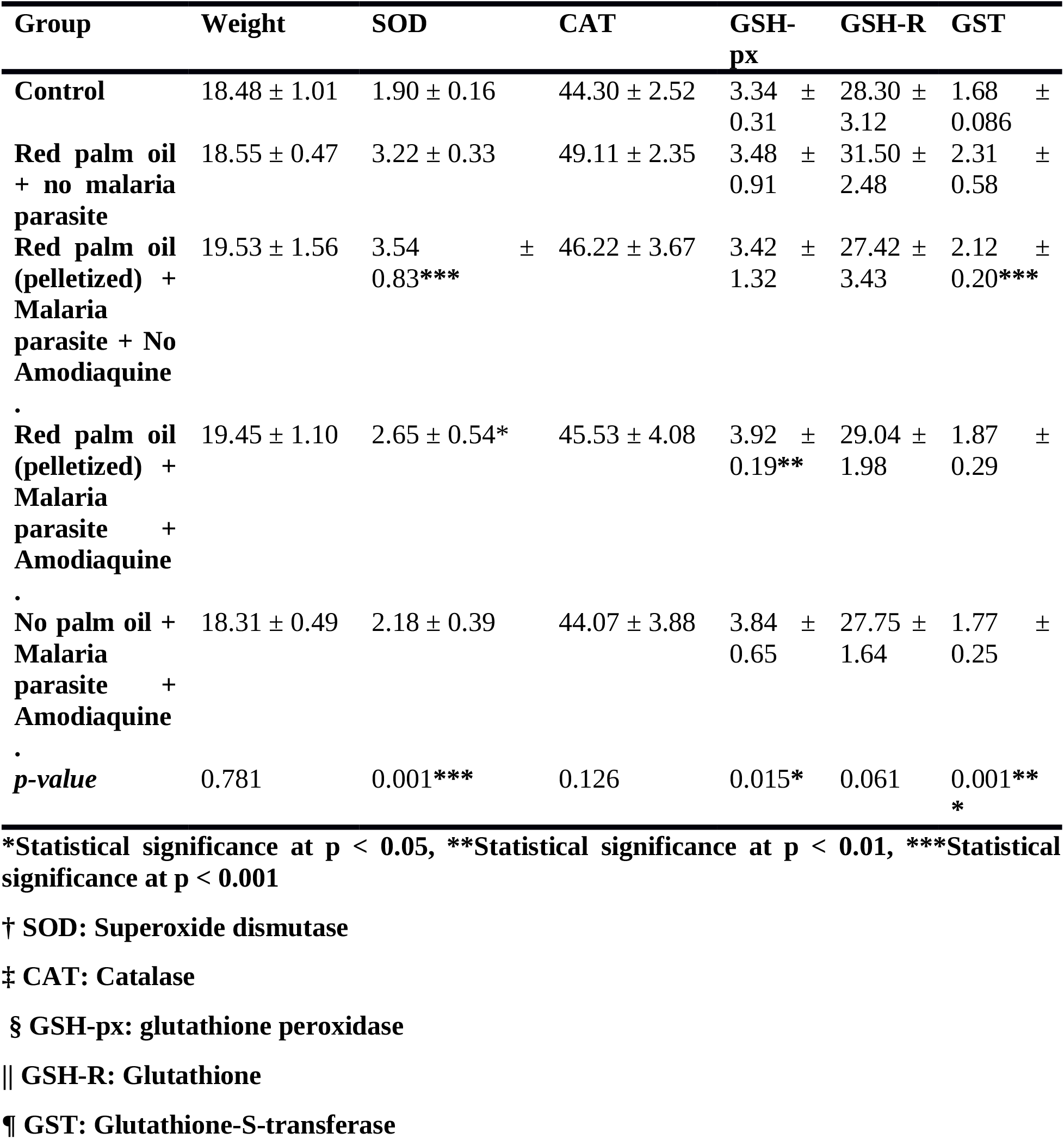
Comparison between the mean ± SD values of Group A and other Groups.

**Table 3:** Comparison between the mean ± SD values of Group B and other Groups.

| Group | Weight | SOD | CAT | GSH-px | GSH-R | GST |
| --- | --- | --- | --- | --- | --- | --- |
| Control | 18.48 ± 1.01 | 1.90 ± 0.16 | 44.30 ± 2.52 | 3.34 ± 0.31 | 28.30 ± 3.12 | 1.68 ± 0.086 |
| Red palm oil + no malaria parasite | 18.55 ± 0.47 | 3.22 ± 0.33 | 49.11 ± 2.35 | 3.48 ± 0.91 | 31.50 ± 2.48 | 2.31 ± 0.58 |
| Red palm oil (pelletized) + Malaria parasite + No Amodiaquine. | 19.53 ± 1.56 | 3.54 ± 0.83 | 46.22 ± 3.67 | 3.42 ± 1.32 | 27.42 ± 3.43 | 2.12 ± 0.20 |
| Red palm oil (pelletized) + Malaria parasite + Amodiaquine. | 19.45 ± 1.10 | 2.65 ± 0.54 | 45.53 ± 4.08 | 3.92 ± 0.19 | 29.04 ± 1.98 | 1.87 ± 0.29 |
| No palm oil + Malaria parasite + Amodiaquine. | 18.31 ± 0.49 | 2.18 ± 0.39*** | 44.07 ± 3.88* | 3.84 ± 0.65 | 27.75 ± 1.64* | 1.77 ± 0.25 |
| *p-value* | 0.893 | 0.003** | 0.042* | 0.772 | 0.031* | 0.787 |

\*Statistical significance at $p < 0.05$ , \*\*Statistical significance at $p < 0.01$ , \*\*\*Statistical significance at $p < 0.001$
† SOD: Superoxide dismutase
‡ CAT: Catalase
§ GSH-px: glutathione peroxidase
||GSH-R: Glutathione
¶GST: Glutathione-S-transferase

**Figure 1:**
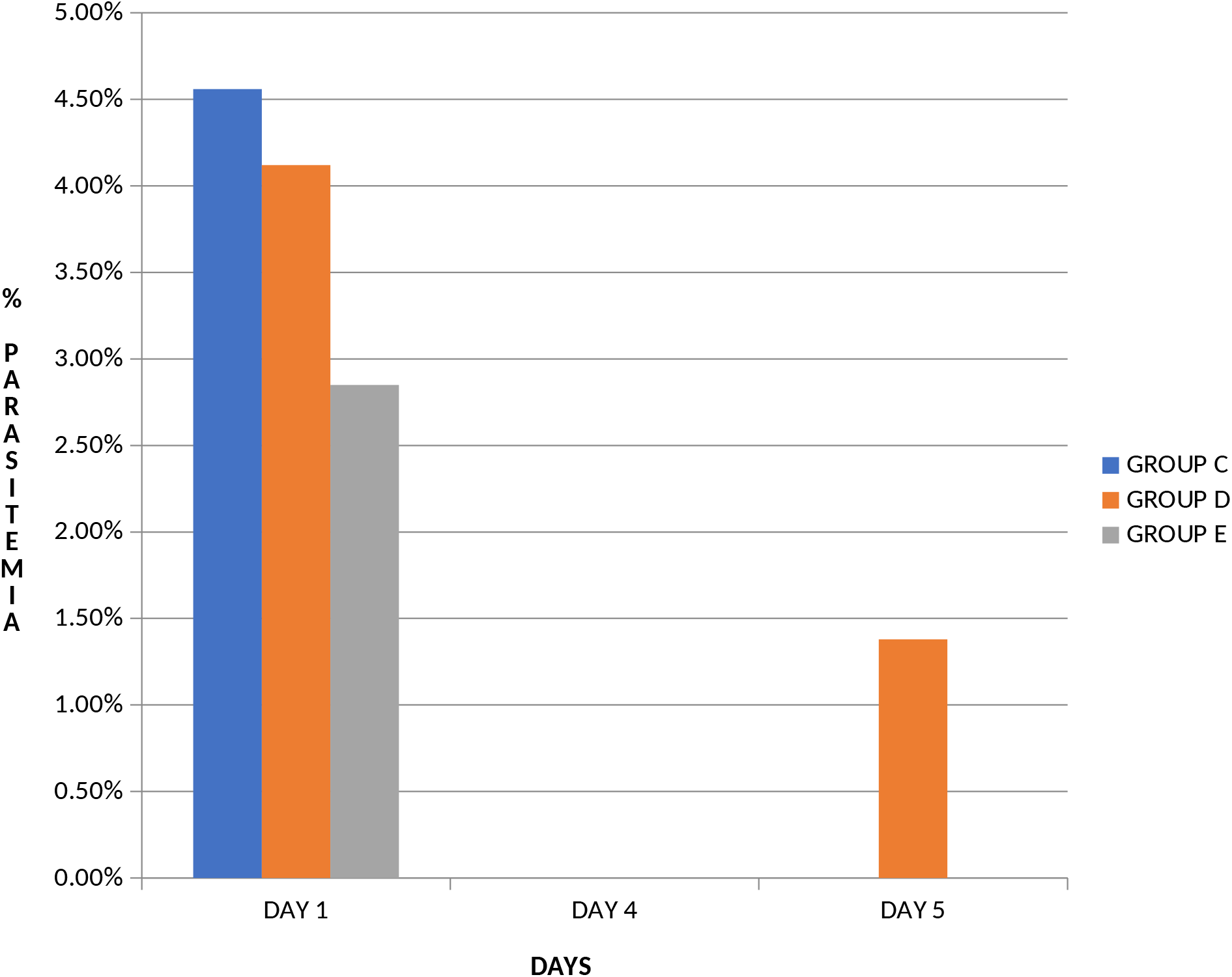
shows the % parasitemia levels in each test group against the number of days.

The study groups comprised of negative and positive controls; Red palm oil (pelletized) + Malaria parasite + No Amodiaquine; red palm oil (pelletized) + Malaria parasite + Amodiaquine; No palm oil + Malaria parasite + Amodiaquine. The mean values of antioxidant enzymes (SOD, catalase and GST) showed statistically significant difference (*p*<0.05) in the positive and negative group [Table 2]. The values of SOD and GST were found to be significantly increased in the group C (with healthy mice fed with red palm oil, infected with malaria parasite without antimalaria drug) when compared to the negative control group (p< 0.001). Similarly, an increased level of SOD and GSH-P_x_ was observed in the group D (mice fed with red palm oil, infected with malaria parasite and treated with amodiaquine) when compared to the negative control group (p<0.05).A significant reduction was observed in the levels of SOD, Catalase and GSH-R in the group E (No palm oil + Malaria parasite + Amodiaquine) when compared to the positive control group (p<0.05) [Table 2].

## Discussion

Previous studies have shown that antioxidants may act directly or indirectly in the neutralization of free radicals [19]. Antioxidant compounds are present both in organisms and in food intake (20). It has been suggested that red palm oil (RPO) could minimize oxidative damage through its potential ability to increase the activity of antioxidant enzymes and it may therefore play a role in the prevention and treatment of oxidative injuries to cells [12]. The study aimed to investigate the relationship between antioxidant properties of red palm oil and *Plasmodium berghei* malaria induced oxidative stress. The mice were fed with diet supplemented with red palm oil, an antioxidant in the proportion of 15ml to 1000g of the feed. Oxidative stress enzymes biomarkers SOD, catalase, GSH-P_x,_ GSH-R and GST were analyzed and compared with each group.

From this study, a statistical significance difference (p<0.05) was observed in table 2 which showed increase in the activities of superoxide dismutase, catalase, and glutathione transferase of liver homogenate samples of mice in Group B (positive control) when compared to Group A (negative control). There was an indication that red palm oil has significantly increased the level of antioxidant enzymes activities in group B (healthy mice fed with red palm oil without malaria parasite) and this is in agreement with some studies that different concentrations of RPO have differential effects to increase the activities of antioxidant enzymes [21,22].

A statistically significant increase was observed in the levels of SOD and GST in samples of mice in group C when compared to A as shown in table 2. This is an indication that the RPO contained antioxidant enzymes which has helped to ameliorate the increase in generation of free radicals generated by malaria infection. This is evident in the level of parasitemia as shown in Figure 1 on day 1 and day 4 after the day of inoculation. This observation is consistent with previous study elsewhere [23].

The level of antioxidant enzymes (SOD and GSH-Px) increased significantly when mice in group D were compared to group A [Table 2]. This is also in agreement with [23] where it was found out that possibility of good and early recovery was discovered when antioxidant supplementation was given along with antimalarial treatment which is an indication of good prognosis. This can also be ascertained by a decrease to zero in the level of parasitemia count of mice in group D from day 1 to day 4 after days of inoculation. However, a recurrent increased parasitemia level was observed on day 5 after inoculation in group D mice. This observation is consistent with a previous study that a possible factor for the recurrence of malaria maybe as a result of increased rate of poor prophylactic effect of amodiaquine [24]. This level of resistance should be strictly monitored in further studies. Therefore, the role of antimalarial drug (amodiaquine) to suppress the level of malaria parasite on a combined therapy action showed no positive effect.

However, it has been suggested that amodiaquine in combination with artesunate is considered to be highly infective against varying species of malaria parasites [25]. In addition, no significant difference was observed in the level of antioxidant enzymes between A and group E. This could be as a result of increased generation of free radicals thereby causing a shift in the oxidant-antioxidant balance which could lead to tissue damage observed in this disease.

It has been observed there was no significant difference in the level of antioxidant enzymes between the positive control group and group C (treated with red palm oil, malaria parasite without amodiaquine) (Table 3). Meanwhile, the combined therapy of both amodiaquine and RPO should be able to boost the level of activities of antioxidant enzymes as to as ameliorate the effect of oxidative stress in malaria when compared to only antioxidant supplemented diet treatment. Some studies have shown that amodiaquine medication can be used in the treated of uncomplicated malaria [26, 27]. In case of severe malaria, there is a high risk of resistance by the parasite and therefore a combined therapy is allowed.

In addition, the levels of antioxidant enzymes activities (SOD, Catalase and GSH-R) showed significant decrease in mice of group E when compared to group B (Table 3). This has proven another poor prophylactic effect of amodiaquine on malaria infection. The significant decrease of activities of antioxidant enzymes can also be due the fact some antimalarial drugs acted by increasing the oxidant stress level due to the mechanisms of actions of most drugs [28].

Using red palm oil as a substitute for the combined drugs showed no statistical difference and this was unable to combat the activities of free radicals generated by the malaria infection leading to oxidative stress. This was evident on day 5 as a result of a recurrence with high severity when compared to day 4 parasitemia levels. This is also in agreement that have some antimalarial drugs acted by increasing the oxidant stress level due to the mechanisms of actions of most drugs [28].

Increase in the antioxidants levels may be due to successful antimalarial activity of the drug alone which may have further enhanced the ameliorative effect in scavenging all the free radicals (ROS) produced by the parasite, drug action and the natural immune response of the host (mice) [29]. In addition, the deficiency of glutathione reductase enzymes leads to increase production of free radicals such as H_2_O_2_, superoxide which are major oxidants in oxidative stress [30].

## Conclusion

In this study, there is observation of significant increase in the activity of antioxidant enzymes after feeding the parasitized mice with red palm oil. Red palm oil can therefore be said to possess antioxidative properties that ameliorate oxidative stress but not totally eliminate it when combined with antimalarial drugs. However, dietary intake of palm oil helps to reduce free radical mediated injury to the tissue thus preventing oxidative stress induced by malaria and any other related factors.

## Acknowledgements

We appreciate the contributions of the entire staff of Institute for Advance Medical Research and Training (IMRAT), College of Medicine, UCH, Ibadan, Oyo State, Nigeria towards the completion of this study.

